# sumSTAAR: a flexible framework for gene-based association studies using GWAS summary statistics

**DOI:** 10.1101/2021.10.25.465680

**Authors:** Nadezhda M. Belonogova, Gulnara R. Svishcheva, Anatoly V. Kirichenko, Yakov A. Tsepilov, Tatiana I. Axenovich

**Affiliations:** Laboratory of Segregation and Recombination Analyses, Institute of Cytology and Genetics, Siberian Branch of the Russian Academy of Sciences, Novosibirsk, Russia; Laboratory of Animal Genetics, Vavilov Institute of General Genetics, the Russian Academy of Sciences, Moscow, Russia; Department of Natural Sciences, Novosibirsk State University, Novosibirsk, Russia

## Abstract

Gene-based association analysis is an effective gene mapping tool. Many gene-based methods have been proposed recently. However, their power depends on the underlying genetic architecture, which is rarely known in complex traits, and so it is likely that a combination of such methods could serve as a universal approach. Several frameworks combining different gene-based methods have been developed. However, they all imply a fixed set of methods, weights and functional annotations. Moreover, most of them use individual phenotypes and genotypes as input data. Here, we introduce sumSTAAR, a framework for gene-based association analysis using summary statistics obtained from genome-wide association studies (GWAS). It is an extended and modified version of STAAR framework proposed by Li and colleagues in 2020. The sumSTAAR framework offers a wider range of gene-based methods to combine. It allows the user to arbitrarily define a set of these methods, weighting functions and probabilities of genetic variants being causal. The methods used in the framework were adapted to analyse genes with large number of SNPs to decrease the running time. The framework includes the polygene pruning procedure to guard against the influence of the strong GWAS signals outside the gene. We also present new improved matrices of correlations between the genotypes of variants within genes. These matrices estimated on a sample of 265,000 individuals are a state-of-the-art replacement of widely used matrices based on the 1000 Genomes Project data.

**AUTHOR SUMMARY:** Gene-based association analysis is an effective gene mapping tool. Quite a few frameworks have been proposed recently for gene-based association analysis using a combination of different methods. However, all of these frameworks have at least one of the disadvantages: they use a fixed set of methods, they cannot use functional annotations, or they use individual phenotypes and genotypes as input data. To overcome these limitations, we propose sumSTAAR, a framework for gene-based association analysis using GWAS summary statistics. Our framework allows the user to arbitrarily define a set of the methods and functional annotations. Moreover, we adopted the methods for the analysis of genes with a large number of SNPs to decrease the running time. The framework includes the polygene pruning procedure to guard against the influence of the strong GWAS signals outside the gene. We also present new improved matrices of correlations between the genotypes of variants within genes, which now allows to include ultra-rare variants in analysis.

## INTRODUCTION

Gene-based association analysis is an effective replacement of genome-wide association analysis (GWAS) in identification of rare genetic variants ^1, 2^. Many gene-based methods have been proposed recently. Their power depends on the underlying genetic architecture that is rarely known in complex traits. Therefore, a combination of such methods could serve as a universal approach.

Among popular combined tests, SKAT-O was the first, for which the distribution of test statistics was analytically described ^3^. For other combined tests, p-values were estimated empirically at the cost of dramatically increased computation time. The task to analytically combine the p-values obtained by different methods has been solved in the aggregated Cauchy test, ACAT-O ^4^. This gave impetus to create a range of frameworks in order to one-by-one calculate a number of gene-based tests and then combine them by ACAT-O ^5–9^. The frameworks differ from one another by the task, input data, methods used, and ways to include functional biological annotations. All these frameworks have a disadvantage: they are not flexible. They use the fixed set of methods, weights and combinations of functional annotations. Moreover, the majority of existing frameworks use individual phenotypes and genotypes as input data. Such data cannot be deposited in open-access databases, and so they are unavailable to a wide range of investigators. Recently, it was demonstrated that all popular methods of gene-based association analysis based on the linear regression models can use summary statistics instead of individual data ^10^. Previously, we presented formulas for the wide range of association tests and implemented them in the sumFREGAT package ^11^.

The framework named STAAR (variant-set test for association using annotation information) stands out among others as a comprehensive and powerful tool that effectively incorporates SNP-weighting by allele frequencies, variant categories and multiple complementary annotations^6^. Here we propose the extended and modified version of the STAAR framework, which we called sumSTAAR. The modification concerns the input data: STAAR uses raw genotypes and phenotypes, and sumSTAAR uses GWAS summary statistics (effect sizes, standard errors, sample sizes etc.). Extension relates to the gene-based association analysis methods used: STAAR uses a fixed set of three methods, and sumSTAAR uses an arbitrary set including up to six methods.

The methods comprised by the sumSTAAR framework are modified in two ways compared with those previously described.^11^ First, they involve a special algorithm for the analysis of large genes with >500 SNPs. Second, they use more effective computational algorithms for matrix operations. An additional empowering feature of the framework is the use of new LD matrices estimated on an extended sample: 265K instead of 503 individuals from the 1000G project. For the first time, due to these high-coverage estimates, it became possible not to discard the large amount of rare variants when analyzing summary statistics with a wide range of powerful gene-based methods. We also present the procedure of polygene pruning to guard against the influence of strong association signals outside the gene on the results of gene-based association analysis ^12^.

## METHODS

### Gene-based association analysis

The sumSTAAR framework combines (a set of) the following methods: burden test (BT), SKAT, SKAT-O, aggregated Cauchy association test (ACAT-V), the tests using functional linear regression model (FLM) and principal component analysis (PCA). Variant-specific weights can be applied to all of these methods. We also introduced the probabilities of genetic variants being causal estimated using different functional annotations in BT, SKAT, SKAT-O, FLM, PCA and ACAT-V. All these modified tests and the parameters of their distributions are presented in *Supplementary Materials*. The sumSTAAR() function (Fig. 1) allows the user to arbitrarily choose a set of tests that differ in method, weighting function and probabilities of genetic variants being causal, calculate these tests, and then combine them using ACAT-O.

**Figure 1.**
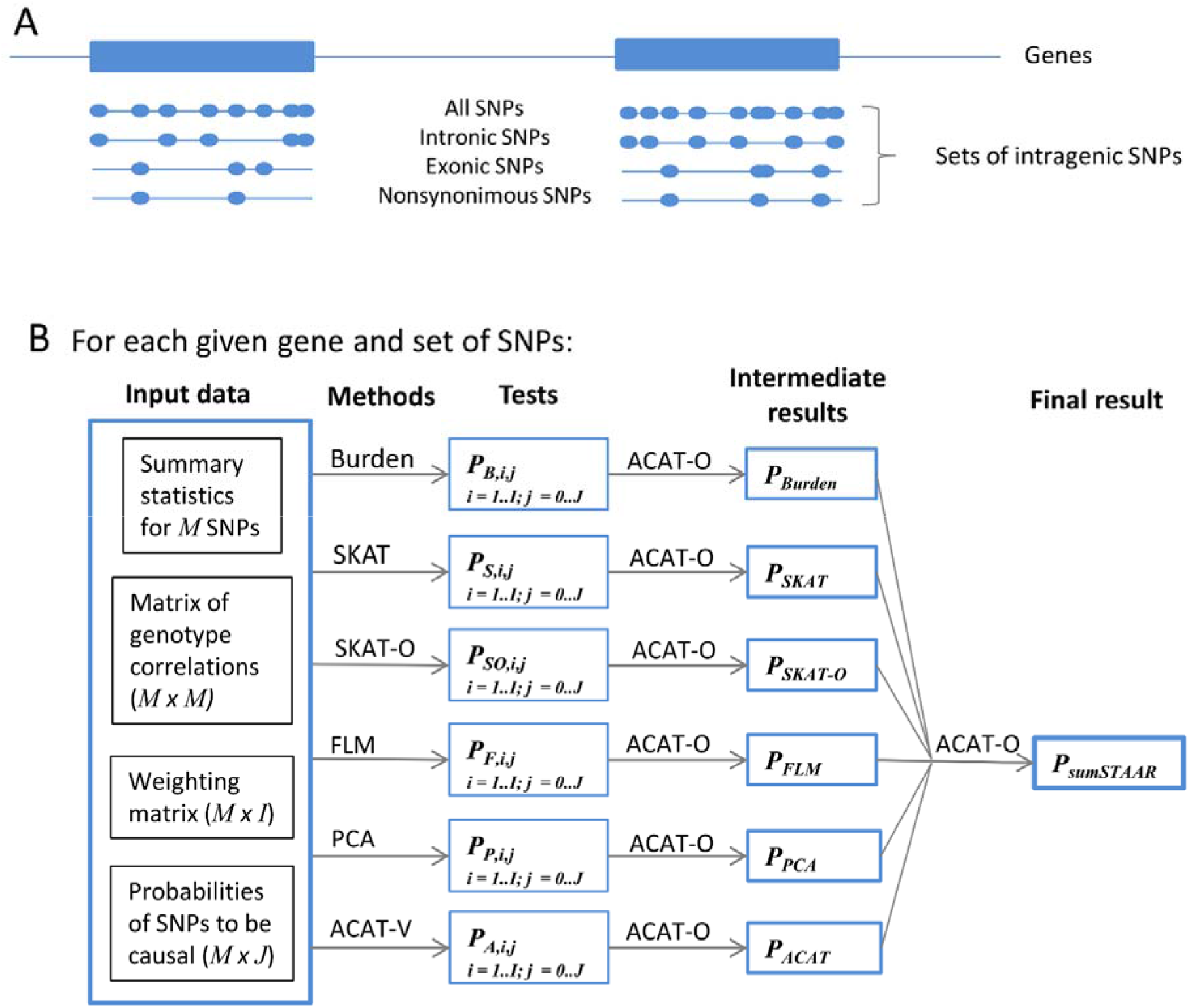
Workflow schematic. (A) Each set of SNPs (all, non-coding, exonic, nonsynonymous and others) is analyzed separately. (B) Input data for sumFREGAT include GWAS summary statistics (p-values and effect sizes), correlations between genotypes calculated using the same or reference sample, the matrix of weighting functions defined by the parameters of the beta distribution, the probabilities of SNPs being causal (e.g., estimated using different functional annotations http://favor.genohub.org/). The list of methods can comprise an arbitrary subset of BT, SKAT, SKAT-O, PCA, FLM, and ACAT-V. For each method, regionbased association analysis is repeatedly performed using different combinations of the weighting functions (*i* ∈ [1, *I*]) and probabilities of SNPs being causal (*j* ∈ [0, *J*]). ACAT-O is used for combining the p-values obtained by each method under different weighting functions and probabilities, and then for combining the results obtained by various methods.

### Analysis of large genes

To decrease the running time for association analysis of genes with a large number of SNPs, we propose the following algorithm. Using thresholding technique, we divide all SNPs within a gene into two groups by p-values, which correspond to their weighted z-scores. Since multiple linear regression models include SNP-specific weights, we form SNP groups taking into account these weights. We used a threshold of 0.8, which was selected empirically (see below). We apply a chosen gene-based test to the group with the smaller weighted p-values and calculate the simple mean weighted p-value for another group. Then we combine p-values obtained for the two groups by ACAT-O. Obviously, this algorithm is an approximation to the chosen gene-based test, however it proves to be very effective for the genes with more than 500 SNPs. We introduced it in SKAT, SKAT-O, PCA and FLM methods.

### Selecting the threshold

To choose the threshold, below which weighted p-values are considered as small when analyzing large genes, we performed an empirical assessment of the approximation on the material of summary statistics for neuroticism from UK Biobank dataset ^13^(for details, see *Supplementary Materials*). We calculated the approximated SKAT statistics using a range of values as threshold (from 0.05 to 0.95) on 2,103 genes having from 500 to 10,000 SNPs. For each threshold value, we measured the total elapsed time and calculated *R*^2^ between the original and approximated SKAT statistics (log10(*p*-values)). Since the approximated SKAT p-values deviated in both directions from original ones, we assessed both deviances using the formula

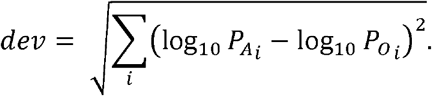

Here *P_Ai_* and *P_Oi_* are the approximation and original p-values for the *i-*th gene, respectively; *dev* for conservativeness and inflation of the approximated test statistics was calculated using *i* ∈ {*P_Ai_* > *P_Oi_*} and *i* ∈ {*P_Ai_* < *P_Oi_*}, respectively.

### LD matrices

The LD matrices of genotype correlations are required as input data for all packages using summary statistics. Such matrices are obtained using reference samples of genotypes, usually from the 1000 Genomes Project. However, due to a small sample size, rare variants are not involved in the calculation of correlation matrices. Using the UK Biobank resource under application #59345, we calculated Pearson correlation coefficients (*r*) between genotypes of variants within gene for 19,426 genes using 265,000 participants of European ancestry from the UK Biobank cohort ^14^ and LDstore software v1.1 ^15^. Only variants with MAF>10^-5^ and imputation quality *r*^2^>0.3 were used for the calculations.

Sociodemographic, physical, lifestyle and health-related characteristics of the UK Biobank cohort have been reported by ^16^. In brief, individuals enrolled in the UK Biobank study were aged 40-69 years; were less likely to be obese, to smoke, to drink alcohol; had fewer self-reported health conditions as compared to the general population. All study participants provided written informed consent, and the study was approved by the North West Multi-Centre for Research Ethics Committee (11/NW/0382). The participants of European ancestry (defined by SNP-based principal component analysis) were randomly selected to provide the GWAS discovery cohort. Genotyping and imputation data were obtained from the UK Biobank March 2018 data release. Genotyping was conducted using the Affymetrix UK BiLEVE and Affymetrix UK Biobank Axiom arrays. Imputation was performed with the IMPUTE4 program (https://jmarchini.org/impute-4/) ^13^ using the Haplotype Reference Consortium (HRC) ^17^ and merged UK10K and 1000 Genomes phase 3 reference panels. Details on the centralized analysis of the genetic data, genotype quality, properties of population structure and relatedness of the genetic data, and efficient phasing and genotype imputation have been reported previously ^13^.

## RESULTS

All three gene-based methods implemented in STAAR (BT, SKAT and ACAT-V) are available in the sumSTAAR framework. We analytically showed the equivalence of these tests between the frameworks (see *Supplementary Materials*) and numerically compared their results obtained in STAAR and sumSTAAR. STAAR implies an omnibus weighting scheme of combining multiple differently weighted tests (see *Supplementary Materials*, Fig. S1). Using summary statistics, we reproduced this scheme in sumSTAAR. The R code to perform simulations and comparisons is available at https://github.com/nbelon/sumSTAAR-vs-STAAR-comparison/blob/main/sumSTAAR.vs.STAAR.R. As can be seen in *Supplementary Materials*, Fig. S2, there is excellent agreement between the results obtained by two packages.

Then, we tested the new algorithm developed for analysis of large genes and picked up the threshold separating the two groups of SNPs in accordance with their weighted p-values. We tried to find a reasonable compromise between approximation accuracy and computation time. We observed the highest correlation between the original and approximated test statistics for threshold values ranging from 0.6 to 0.8 (Fig.2A). There was no prominent change in total elapsed time among these runs. However, the approximated test proved to be more conservative and inflation less frequent with increasing threshold values (Fig.2B). Therefore, to prevent an increase in false positive rate, we selected the threshold of the weighted p-value = 0.8 to separate the SNPs on two groups.

**Figure 2.**
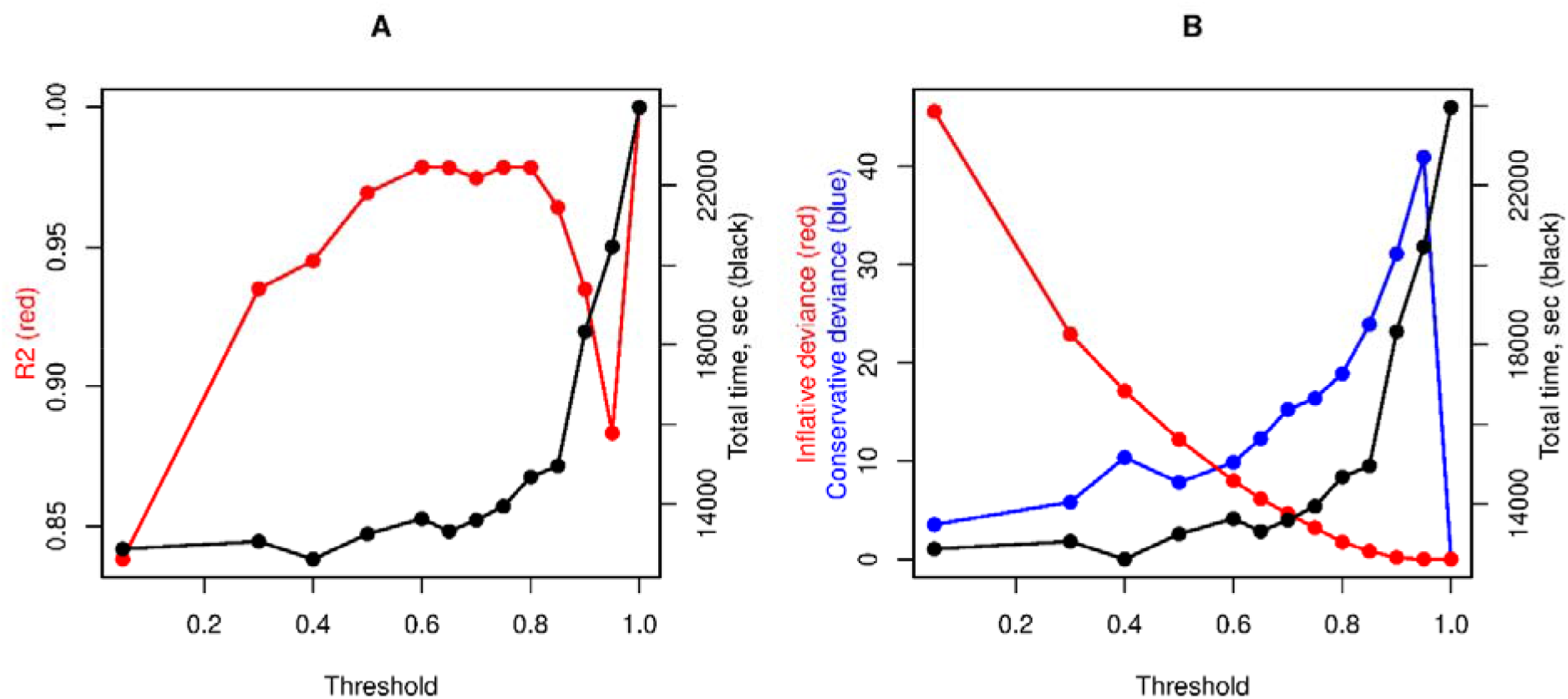
Determination coefficient and deviances of approximated SKAT statistic related to the threshold value. (A) Determination coefficient (*R*^2^) between −log10(P value) of original and approximated tests shown in red. (B) Deviances indicating inflation and conservativeness of approximated test statistics compared with original shown in red and blue, respectively.

Using the neuroticism summary statistics, we estimated the accuracy and efficiency of the modified methods implemented in the new version compared to the version of sumFREGAT without modifications. Figure 3 shows a good agreement between the p-values obtained by two packages and a decrease in the running time when using the new modified version of the package. For SKAT, SKAT-O, PCA and FLM methods, the running time was decreased by 2.4, 10.5, 3.4 and 2.6 times, respectively. The most prominent effect was shown for the most popular SKAT-O method.

**Figure 3.**
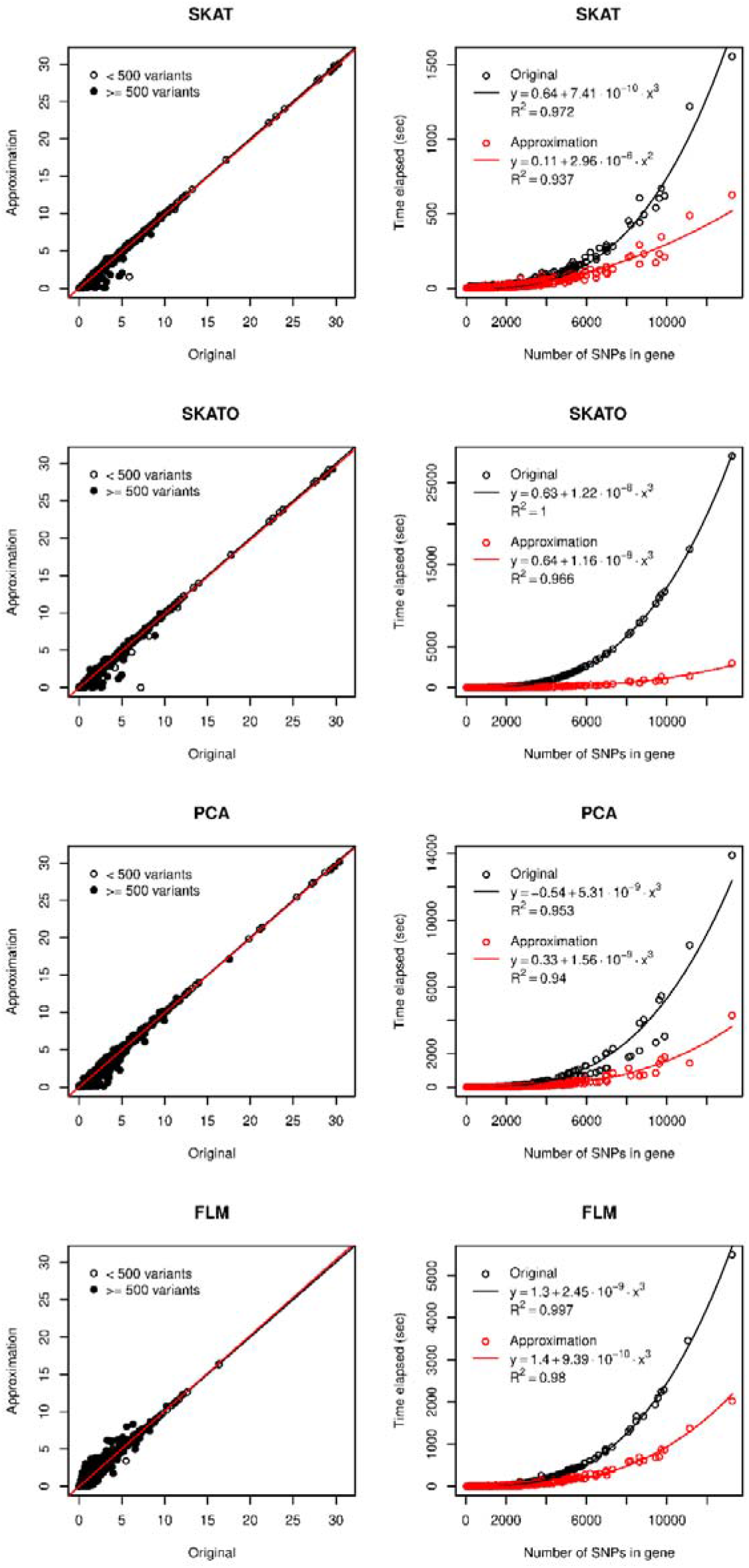
Accuracy and running time of four gene-based methods for association analysis under approximation. Each point represents a gene: 7,990 genes for FLM (genes that passed collinearity filter for 25 basis functions, see *Supplementary Materials* for details) and 17,975 genes for other methods. Left panels show −logl0(P values), red lines are regression lines and black lines represent one-to-one correspondence. On the right panels, lines represent the best-fitted polynomial functions.

Within the framework, we introduce the new LD matrices for 19,426 genes estimated using genotypes of 265K participants of UK Biobank project. The matrices contain information about 21,155,091 SNPs, with 17,142,006 (81%) of them having MAF < 0.01. For comparison, the widely used matrices of SNP-SNP correlations estimated on 503 European participants of 1000G project include 4,544,901 SNPs, with only 707,862 (16%) of them having MAF < 0.01. The UK Biobank matrices, therefore, provide 4.65 times higher SNP coverage and 24 times higher coverage for rare variants.

For the polygene pruning procedure, we now publish an R-script to perform it step-by-step.

## DISCUSSION

We developed a new framework for gene-based association analysis using summary statistics. This framework can include an arbitrary sets of methods for gene-based association analysis, weighting functions and functional annotations used for the estimation of SNP probability being causal. Many of the methods used in the framework were adapted to the genes with large number of SNPs. This allows us to increase the computation speed of different methods by 2.4 – 10.5 times.

We compared STAAR and sumSTAAR and demonstrated the strong agreement of the results obtained by BT, SKAT, ACAT-V and their ACAT-O-based combinations. Our sumSTAAR framework, however, provides an opportunity to expand the range of methods with the fixed-effects models (PCA and FLM). In addition, SKAT-O can be used instead of ACAT-O to combine the BT and SKAT. Being more computationally intensive, SKAT-O nevertheless provides an optimal, kernel-based combination of the methods and higher statistical power compared with ACAT-O.

We introduced the new LD matrices with high-coverage estimates of SNP-SNP correlations for human genes. Due to these matrices, it became possible to add the large amount of rare variants when analyzing summary statistics with a wide range of gene-based methods. The matrices can be used not only in our software, but also in any program for genetic analysis using summary statistics.

The sumSTAAR framework suggests using the polygene pruning procedure to guard against the influence of the strong GWAS signals outside the gene. It has been shown that a substantial share of gene-based association signals is inflated by these GWAS signals ^6, 12^. To guard against this influence, the conditional GWAS summary statistics calculated using, for example, the GCTA-COJO package ^18^ can be used in sumFREGAT as input data. However, to calculate the conditional statistics, this type of analysis relies on the simple multiple regression with all the attendant limitations. For example, conditional SNPs should be in complete linkage equilibrium with each other and their number, therefore, cannot be large. The procedure called “polygene pruning” ^12^ represents an alternative way to reduce the effect of strong GWAS signals outside the gene. Polygene pruning results in exclusion of some variants within the gene being in LD with outside GWAS-identified variants from gene-based analysis. In essence, this procedure is analogous to the extreme weighting of within-gene SNPs based on their LD with outer GWAS signals. Polygene pruning way can be preferable when the set of within-gene variants is large or includes rare variants. Moreover, the classical conditional analysis is impossible to perform when genotypes of top GWAS signals are not available, while correlation structure sufficient for polygene pruning can be shared more easily.

To our knowledge, sumSTAAR is the most flexible and one of the most comprehensive frameworks that allow researchers to perform state-of-the-art gene-based analyses using GWAS summary statistics.

## DATA AND SOFTWARE AVAILABILITY

sumFREGAT: https://CRAN.R-project.org/package=sumFREGAT

LD matrices based on UKBB sample: https://mga.bionet.nsc.ru/sumfregat/ukbb/

Polygene pruning: https://github.com/nbelon/Polygene_pruning

GWAS summary statistics used: https://ctg.cncr.nl/documents/p1651/sumstats_neuro_sum_ctg_format.txt.gz

## AUTHOR CONTRIBUTIONS

Conceptualization: GRS TIA YAT. Investigation: AVK GRS NMB. Methodology: GRS NMB. Resources: AVK YAT. Software: GRS NMB. Supervision: TIA YAT. Visualization: NMB. Writing – Original Draft Preparation: TIA. Writing – Review & Editing: GRS NMB TIA YAT.

