## Supplementary material for "sumSTAAR: a flexible framework for gene-based association studies using GWAS summary statistics": Suppl_Meth_Bioarchive.docx

**Introducing the probabilities of genetic variants being causal in the different methods**

A great number of summary SNP-level statistics such as z-scores (*z*), p-values (*p*) and effect sizes (*β*) are now available for different traits and diseases in open-access databases (Pasaniuc and Price, 2017). By definition, for each genetic variant, z-score follows the standard normal distribution *N*(0,1) under the null hypothesis of no association between the variant and the trait, and can be easily calculated from another GWAS summary statistics, namely the p-value and *β*: the absolute value of a z-score is defined by the p-value as: |*z*| = Φ^-1^(1-*p*/2), where Φ is a standard normal cumulative distribution function, and the z-score sign is defined by the sign of the corresponding *β*.

For a sample of unrelated individuals, a vector of z-scores, *Z*, obtained one by one for each of *M* SNPs in the gene is defined as:

$$Z=\frac{V^{-1}\tilde{G}^{T}\tilde{y}}{\sqrt{N}\sigma_{y}}.$$

Here $\tilde{G}$ and $\tilde{y}$ denote the centered values of an (*N×M*) matrix of genotypes and an (*N×*1) vector of phenotypes, respectively; *V* is an (*M×M*) diagonal matrix of the square roots of genotypic variances defined via $\sqrt{se\left( \beta\right)}$ or $\sqrt{MAF\left( 1-MAF \right)}$ under Hardy-Weinberg equilibrium, where *se*() denotes standard errors, and *MAF* is minor allelic frequencies; *N* is the sample size and *σ_y_* is the phenotypic variance.

For *M* genetic variants, *Z* follows the multivariate normal distribution *N*(0,*U*) under the null hypothesis of no association between the genetic variants and the trait, where *U* is calculated as an (*M*×*M*) matrix of correlations between the genotypes of these variants (Conneely and Boehnke, 2007):

$$U=\frac{V^{-1}\tilde{G}^{T}\tilde{G}V^{-1}}{N}.$$

The *U* matrix can be estimated using a reference sample of genotypes originating from the same population.

In the methods implemented in sumFREGAT and using the multiple linear regression models, the test statistics and their distributions are presented in terms of *Z* and *U* that can be weighted in a certain way.

For Burden test (BT) and SKAT based on the random effects models, test statistics are:

$$\begin{matrix} Q_{BT,i,j}=\left( Z^{T}VW_{i}\Pi_{j}e \right)^{2}, \\ Q_{SKAT,i,j}=Z^{T}VW_{i}\Pi_{j}W_{i}VZ, \end{matrix} (1)$$

where *W_i_* is a diagonal matrix of SNP weights defined via the beta density function of MAFs with the *i*-th set of parameters; *Π_j_* is a diagonal matrix of the probabilities of genetic variants being causal defined by the *j*-th functional annotation, and *e* is the vector of units.

Under the null hypothesis, *Q_BT_* follows a scaled χ^2^ distribution with one degree of freedom, $\lambda\chi_{\mathrm{df}=1}^{2}$, where *λ* $=e^{T}\Pi_{j}W_{i}VUVW_{i}\Pi_{j}e$, and *Q_SKAT_* follows a weighted sum of χ^2^_df=1_ distributions, $\sum\lambda\chi_{\mathrm{df}=1}^{2}$, where *λ* are eigenvalues of $W_{i}VUVW_{i}\Pi_{j}$*.*

Another kernel-based method presented in sumFREGAT is SKAT-O (Lee, et al., 2012; Svishcheva, 2019; Svishcheva, et al., 2019). Since this method is a linear combination of BT and SKAT, *Q_SKATO_* follows the weighted sum of χ^2^_df=1_ distributions, where weights are the eigenvalues of the kernel matrix*.* For SKAT-O, the probabilities of genetic variants being causal (*Π_j_*) can be introduced into test statistics and eigenvalues of the kernel matrix as follows:

$$\begin{matrix} Q_{SKATO,i,j}=\rho\left( Z^{T}VW_{i}\Pi_{j}e \right)^{2}+\left( 1-\rho\right)Z^{T}VW_{i}\Pi_{j}W_{i}VZ \\ \lambda=eigen\left( \rho\left( U^{\frac{1}{2}}VW_{i}\Pi_{j}ee^{T}\Pi_{j}W_{i}VU^{\frac{1}{2}} \right)+\left( 1-\rho\right)\left( U^{\frac{1}{2}}VW_{i}\Pi_{j}W_{i}VU^{\frac{1}{2}} \right) \right). \end{matrix}$$

where *ρ* is the coefficient of the linear combination of tests.

The methods based on multiple linear regression models with fixed genetic effects use the F-test statistic to test the null hypothesis H_0_: *β* = 0 against H_1_: *β* ≠ 0. For the complete multiple linear regression model (MLR), the F-test statistic for a region with *M* genetic variants is calculated by (Chow, 1960):

$$F=\frac{R^{2}\left( N-M-1 \right)}{\left( 1-R^{2} \right)M},$$

where

$$R^{2}=\frac{Z^{T}U^{-1}Z}{N}.$$

The p-value for MLR is defined by an F-distribution with *M* and *N-M-*1 as the numbers of degrees of freedom. As can be seen, MLR requires the inverse of the correlation matrix *U*. If the columns of *U* are collinear or nearly collinear, numerical instability and potentially inflated regression coefficients may appear. On average, collinearity in *U* substantially depends on SNP density, i.e. the number of SNPs analyzed. New dense correlation matrices proposed in this study provide 4.65 times higher SNP coverage and, therefore, higher collinearity in *U*. Taking into account the sensitivity of MLR to the increased collinearity, we did not include MLR in our updated version of sumFREGAT.

In contrast to MLR, the PCA (Wang and Abbott, 2008) and FLM (Belonogova, et al., 2018; Fan, et al., 2013; Svishcheva, et al., 2015) methods use incomplete (truncated) linear regression models with fixed effects. Both methods serve to reduce the number of regression predictors by generating an orthogonal basis set of *K* elements. For PCA, these elements are the first *K* principal components containing a large proportion of information about genotype data. For FLM, these elements are the *K* basis functions, by which the genotypes and their effects can be presented as continuous functions. For incomplete regression models, *R*^2^ is calculated as

$$R^{2}=\frac{Z^{T}{VW_{i}C\left( C^{T}W_{i}VUVW_{i}C \right)}^{-1}C^{T}W_{i}VZ}{N}, \left( 4 \right)$$

where *C* is the (*M*×*K*) matrix specified for each gene-based method (Belonogova, et al., 2018; Svishcheva, et al., 2015). For FLM, each element of *C*, *C_mk_*, represents the value of the *k*-th basis function calculated at the *m*-th genetic variant position. For PCA, *C* is given as a truncated matrix of right singular vectors obtained from the singular value decomposition of the weighted genotype matrix. Truncating is achieved by considering only the first *K* largest squared singular values that account for 80-90% of the total genotype variance observed in the genomic region (Gauderman, et al., 2007; Wang and Abbott, 2008). For methods using incomplete multiple linear regression models, the value of *M* in the F-test numbers of degrees of freedom is replaced by

$$M'=rank\left( C^{T}W_{i}VUVW_{i}C \right).$$

The probabilities of genetic variants being causal (*Π_j_*) are introduced in Expression (4) as follows:

$$R^{2}=\frac{Z^{T}{VW_{i}\Pi_{j}C\left( C^{T}\Pi_{j}W_{i}VUVW_{i}\Pi_{j}C \right)}^{-1}C^{T}\Pi_{j}W_{i}VZ}{N}$$

and

$$M'=rank\left( C^{T}\Pi_{j}W_{i}VUVW_{i}\Pi_{j}C \right).$$

PCA is a robust method, because multicollinearity is completely eliminated by orthogonal conversion of correlated predictors to principal components (PCs) that are linearly uncorrelated. PCA always ensures *Mʹ* = *K*.

In contrast, the FLM results are not always stable because FLM minimizes but does not eliminate multicollinearity. The loss of orthogonality still may occur in the matrices of basis functions calculated for some genes. To guard against this problem, we introduced a filter to ensure that matrix to be inverted in FLM is full rank (*Mʹ* = *K*). Due to this filter, FLM is applied to a restricted set of genes.

One more method introduced in sumFREGAT is an aggregated Cauchy association test (ACAT) proposed by (Liu, et al., 2019) and modified by (Li, et al., 2020). The gene-based analysis test is defined as:

$$T_{ACAT-V_{i,j}}=\frac{1}{W_{sum}}\sum_{m=1}^{M} {\pi_{j}}_{m}{w_{i}}_{m}^{2}\sigma_{g_{m}}^{2}\tan\left\{ \left( 0.5-p_{m} \right)\pi\right\},$$

were ${\pi_{j}}_{m}$ is the probability of *m*-th variant being causal defined by the *j*-th functional annotation; ${w_{i}}_{m}$ is the weight of *m*-th variant defined by the beta density function of MAF with the *i*-th set of parameters; here $\sigma_{g_{m}}^{2}$ is introduced to make the total weights in ACAT-V comparable with those in methods using linear regression models, $\sigma_{g_{m}}=V_{mm}$; and $W_{sum}=\sum_{m=1}^{M} {\pi_{j}}_{m}{w_{i}}_{m}^{2}\sigma_{g_{m}}^{2}$.

To combine *L* different methods, ACAT is used without any weighting (ACAT-O):

$$T_{ACAT-O}=\frac{1}{L}\sum_{l=1}^{L} \tan\left\{ \left( 0.5-p_{l} \right)\pi\right\}.$$

The p-values for ACAT-V and ACAT-O tests are calculated as:

$$p_{ACAT}= 0.5-\frac{arctg\left( T_{ACAT} \right)}{\pi}.$$

**Equivalence of methods implemented in the sumFREGAT and STAAR packages**

The formulas for gene-based association test statistics proposed by (Li, et al., 2020) are written in terms of *s* but not *z*-score statistics. It is easy to see that

$$S=VZ. (2)$$

Formula (2) follows from the equality $G^{T}\tilde{y}=\tilde{G}^{T}\tilde{y}$ explained by the idempotent property of the centering projection *P* matrix: *PP=P*, where *P* = *I_N_* – *X*(*X^T^X*)^-1^*X^T^*, and *X* is the matrix of covariates taking into account the intercept. Since $\tilde{y}$ can be expressed as $\tilde{y}=Py$,

$G^{T}\tilde{y}=G^{T}Py=G^{T}PPy=\tilde{G}^{T}\tilde{y}$.

To rewrite the formulas for gene-based association tests proposed by (Li, et al., 2020) in terms of z-scores, we present these formulas in matrix form:

$$\begin{matrix} Q_{Burden,i,j}=\left( \sum_{m=1}^{M} \pi_{jm}w_{im}S_{m} \right)^{2}=S^{T}W_{i}\left( \Pi_{j}ee^{T}\Pi_{j} \right)W_{i}S, \\ Q_{SKAT,i,j}=\sum_{m=1}^{M} \pi_{jm}w_{im}^{2}S_{m}^{2}=S^{T}W_{i}\left( \Pi_{j} \right)W_{i}S. \end{matrix} (3)$$

Substituting Expression (2) into Expressions (3) gives

$Q_{BT,i,j}={(Z^{T}VW_{i}\Pi_{j}e)}^{2}$,

$Q_{SKAT,i,j}=Z^{T}VW_{i}\Pi_{j}W_{i}VZ$.

Therefore, Formulas (3) proposed by (Li, et al., 2020) can be presented in terms of summary z-score statistics and the *U* matrix*.* The obtained formulas are equivalent to Formulas (1) implemented in sumFREGAT.

We tested the equivalence of the results obtained by STAAR and sumFREGAT. The STAAR procedure suggests using a combination of three gene-based tests (Burden test, SKAT and ACAT) and two sets of Beta distribution parameters: (1, 1) and (1, 25). For each test and each set of parameters, STAAR calculates 11 individual tests: the original test and 10 tests weighted for each of 10 annotations. The resulting values are combined using the Cauchy method into six STAAR tests (see Fig. 1). P-values from all original (A0) tests are also combined into the ACAT_O p-value. All individual test p-values are combined into the STAAR_O p-value.

Using summary statistics, we reproduced this procedure in sumFREGAT. With the Example data and code from STAAR, we simulated phenotypes and the probabilities of variants being causal for 10,000 individuals and a 5,000-bp region randomly selected on a chromosome 1,000,000 bp in size. The number of SNPs in the region typically fell in the range from 120 to 170, all SNPs having MAF ≤ 0.05. With the individual genotype and phenotype data, we performed all individual and combined tests using the original STAAR procedure. We also calculated z-scores and effect sizes for each variant and the matrix of genotype correlations *U*.

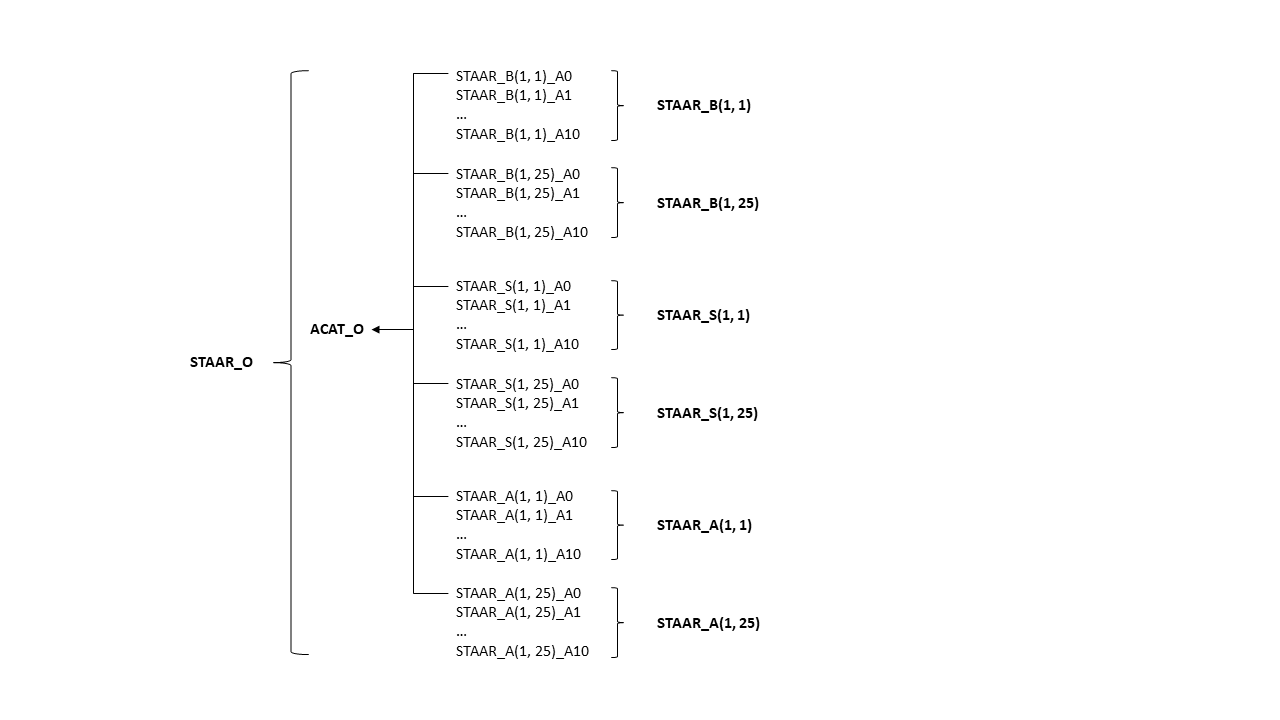

**Fig. 1. Tests performed within the STAAR procedure.** Combined tests are shown in bold.

Next, we used these data as inputs in our sumSTAAR() function of the sumFREGAT package to obtain the same p-values on the summary statistics. R code to perform simulations and comparisons is available at https://github.com/nbelon/sumSTAAR-vs-STAAR-comparison/blob/main/sumSTAAR.vs.STAAR.R. As can be seen in Fig. 2, there is excellent agreement between the results obtained by two packages.

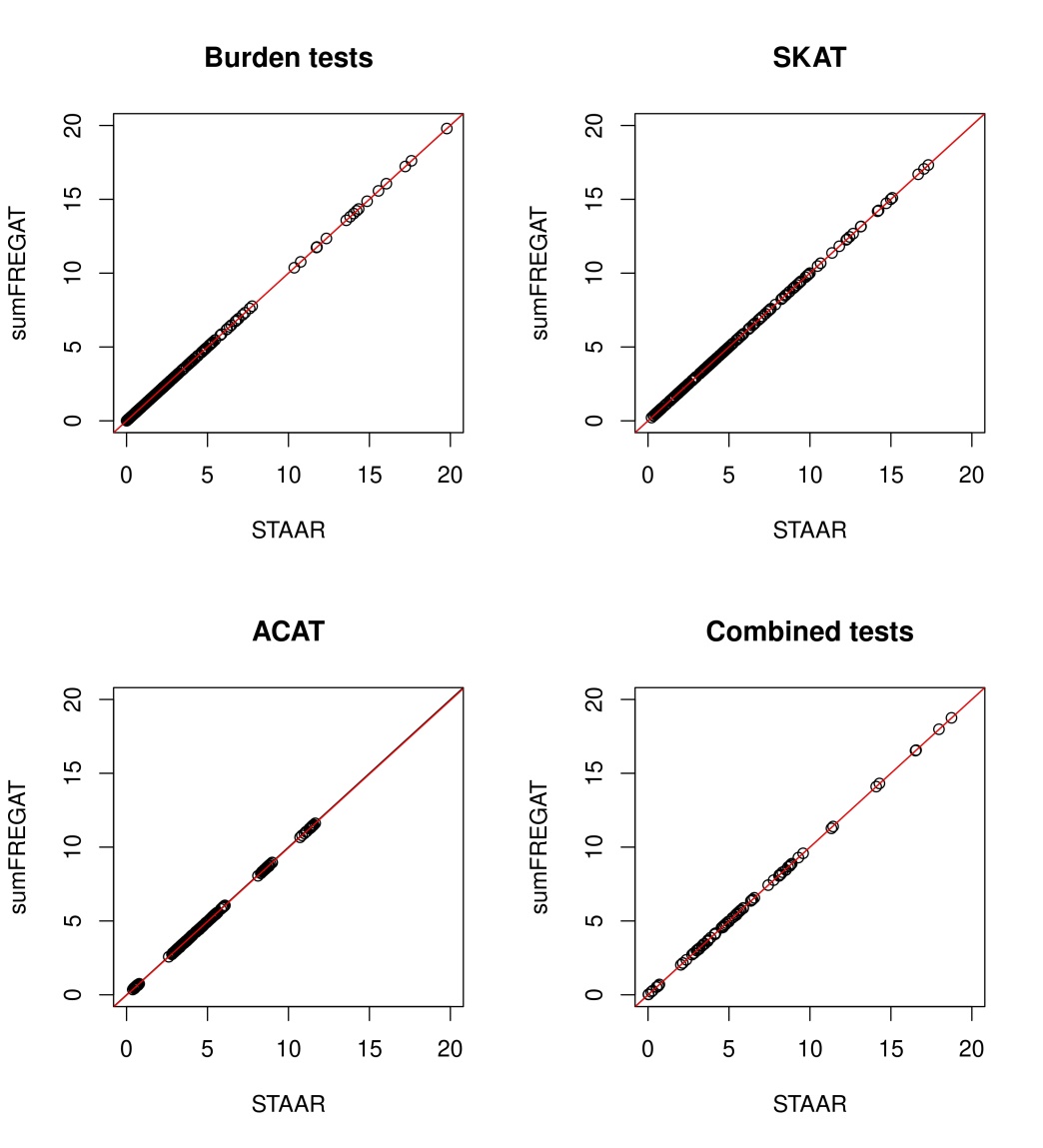

**Fig. 2**. **Comparison of the results obtained by the STAAR and sumFREGAT packages**. Negative log_10_(p-value) were calculated in 10 simulations. The first three panels show the results for individual gene-based tests (Burden test, SKAT and ACAT) with two sets of parameters for the Beta distribution and 11 variants of annotation weighting. The last panel presents –log_10_(p-values) for all combined tests. The regression line is shown in red (overlaps the line of one-to-one correspondence).

### Summary statistics

For testing the modified version of sumFREGAT package, we used the freely available summary statistics (p-values, betas) for neuroticism (<https://ctg.cncr.nl/software/summary_statistics>) obtained in the UK Biobank project (Bycroft, et al., 2018; Sudlow, et al., 2015). These summary statistics for neuroticism were obtained for 10,847,151 SNPs from a sample of 380,506 individuals. SNPs with MAF < 0.001 and INFO < 0.8 were excluded. Neuroticism levels were measured using the Eysenck Personality Questionnaire, Revised Short Form (Eysenck, et al., 1985), consisting of 12 dichotomous items (0, 1). The quantitative trait was defined as a sum of 12 items (for details, see (Nagel, et al., 2018)).
